# Implementing an algorithm for controlling for female MC phase for clinical neuroscience

**DOI:** 10.1101/2020.02.29.971275

**Authors:** Amy A. Herrold, Virginia T. Gallagher, Yufen Chen, Jeffery Majaanes, Natalie Kramer, Brian Vesci, Danielle Colegrove, James Reilly, Leanne McCloskey, Hans Breiter

## Abstract

**Background:** Recent research suggests that hormones and/or menstrual cycle (MC) phase at time of study may significantly affect clinical neuroscience outcomes of interest. Prior research puts forth sound methods for characterizing MC phase in women not using hormonal contraceptives (HC). However, an estimated 40% of premenopausal women in the United States use some form of HC. We developed an algorithm for characterizing hormone levels and MC phase among women both using and not using HC. We have employed this algorithm among female collegiate athletes post-mild traumatic brain injury (mTBI) as female athletes are understudied in the sports mTBI field, MC phase may have important effects on mTBI-related outcomes and controlling for MC phase has not been employed in this population.

**Methods:** Thirty female collegiate athletes were studied. Fifteen incurred a mTBI 3-10 days prior to assessment and sixteen were age, ethnicity, and menstrual cycle (MC) phase matched to the injured athletes. MC matching was conducted with retrospective and prospective self-report MC tracking, self-report of HC use, and serum progesterone testing.

**Results:** 53% (16/30) of females were on HC and 47% (14/30) were not. Of the non-HC users, seven female pairs were in the follicular and one was in the luteal phase. Among the non-HC users, there was 50% agreement in MC phase identification between self-report and progesterone measures and a κ = 0.138. Of the HC users, eight were in the inactive and six were in the active phase of their medication. Among the HC users, average progesterone levels indicated medication compliance (.58ng/mL).

**Conclusions:** This study provides important methodology and proof-of-concept that MC phase can be used as a control variable for time-sensitive prospective clinical neuroscience studies including those involving brain injury. When studying females, it is important to properly examine and control for the sex-specific factor of MC phase in order to have a full understanding of brain behavior relationships in translational research.

## Introduction

Policy put forth by the National Institutes of Health mandates that females be included in all NIH-funded clinical research. When studying females and evaluating sex differences in clinical research, it is important to evaluate the effects of hormone levels and menstrual cycle (MC) phase among females as recent research suggests that hormones and/or menstrual cycle phase at time of study may significantly affect outcomes of interest. Failure to consider the impact of female-specific factors (e.g., MC phase) can also lead to misinterpretation of clinical and potential research biomarkers. For example, cardiometabolic biomarkers fluctuate across the menstrual cycle resulting in differences in classification of cardiovascular risk^1^.

Recent research demonstrates that mild traumatic brain injury (mTBI) results in brain changes, as well as, changes in clinical symptoms, including stress and anxiety^2–4^. However, this research has been conducted almost entirely among males^5^. Studying female athletes is crucial because evidence suggests adolescent females are more vulnerable to injury, experience a greater number of injuries and longer recovery from mTBI relative to males^6–9^. The limited mTBI studies that have included females, have not examined or controlled for female-specific factors such as contraceptive use, hormonal fluctuations, or menstrual cycle (MC) phase at time of injury or assessment^5,10–12^.

We recently reported that collegiate female athletes on hormonal contraceptives had significantly lower post-concussive symptom severity than females not on hormonal contraceptives^6^. Others have shown that menstrual cycle phase and contraceptive use at time of mTBI differentially affects post-concussive symptoms and quality of life one-month post-injury^13^. However, some prior research even suggests excluding females using hormonal contraception^14^ despite current estimates that approximately 82% (43,849 females) of premenopausal, sexually experienced women in the United States have ever used oral contraceptives and approximately 34% (18,074 females) have ever used other hormonal methods of contraception (e.g., NuvaRing®, contraceptive patch)^15^. Thus, excluding females using hormonal contraception is a problematic approach because it limits generalizability. Prior research puts forth sound methods for characterizing menstrual cycle phase in women not using hormonal contraceptives^14^. However, this methodology is not as well characterized for those using hormonal contraceptives.

In this paper, we propose methodology for characterizing hormone levels and menstrual cycle phase among women using hormonal contraceptives (e.g., oral contraceptive pill, Nuva Ring, intrauterine devices) and women not using hormonal contraceptives using both retrospective menstrual cycle tracking, prospective menstrual cycle tracking, and serum progesterone testing. Thus, standard self-report methodology was confirmed and calibrated against hormone levels. Methodology is also presented to match healthy controls to clinical participants on similar hormone levels and menstrual cycle phase to ensure adequate control of these factors in research design. Our goal is to put forth methodology that will enable research studies to feasibly and reliably capture these important factors in clinical research.

## Methods

All research procedures were conducted under an Institutional Review Board-approved protocol.

### Participants

Northwestern University female club athletes between ages 18 and 25 years were recruited for this study. Exclusion criteria: lifetime diagnosis of a psychotic disorder, a first-degree relative with a psychotic disorder, a history of a seizure disorder, and moderate to severe traumatic brain injury. Athletes were excluded from the female athlete control (CON) group if she sustained a concussion within 6 months prior to study participation. An additional component of the study (not discussed here) was identifying neuroimaging markers of mTBI; as such, participants were excluded from the study for contraindications for magnetic resonance imaging (e.g., metal in the body, pregnancy).

Female club athletes diagnosed with mTBI by a physician were recruited to participate in the study, with assessment occurring within 3-10 days following injury. Demographically matched, non-injured, female control athletes were recruited after the respective mTBI athlete completed the protocol.

### mTBI-to-Control Athlete Matching

A control pool of non-collision athletes was created via recruitment at collegiate club athlete events. Once we studied a mTBI athlete, this allowed us to identify a potential control based on age, handedness, ethnicity, and contraceptive use and type. Participants completed a screening form that included the Menstrual History questionnaire from the PhenX Toolkit^16^ to establish whether MCs are typically regular (within 8 days each month) and the approximate length of the cycle (i.e., from the beginning of bleeding of one menstrual period to the beginning of bleeding of the next period). Further, participants indicated current use of hormonal contraceptive (i.e. oral contraceptive pill (OCP), intrauterine device (IUD), contraceptive patch, contraceptive shot, or none). Participants were also asked to provide the start-dates of their three most recent MCs (i.e. the first day of bleeding for the last 3 cycles). Based on this information, participants were assigned to groups: non-hormonal contraceptive users with regular cycles, non-hormonal contraceptive users with irregular cycles, or hormonal contraceptive users. When prospective MC tracking was feasible (i.e. for participants who completed the screening form at least 30 days prior to participation), participants received a prompt via a REDCap generated email to provide the start date of their most recent MC. E-mail prompts were sent based on reported cycle length; for example, if the participant indicated a cycle length of 28-32 days in the screening form, an email prompt was sent approximately every 30 days to provide the first day of their most recent cycle.

### Characterizing MC-phase in non-hormonal contraceptive users

To characterize MC phase at the time of the study visit among female participants not using hormonal contraceptives with regular cycles, we first estimated MC phase by using retrospective and, when available, prospective MC tracking data. Participants in days 1-7 of MC at time of study were estimated to be in the follicular phase at the time of study (Fig. 1). Day 1 is indicated by the first day of self-reported bleeding. Participants in day ≥20 of MC at the time of the study visit were estimated to be in the luteal phase. Participants in days 8-19 of MC at the time of the study visit were not estimated in either follicular or luteal phase due to individual variability in cycles. Serum progesterone levels were collected on the day of study. Regardless of estimated MC-phase based on calendar tracking, participants’ MC phase was classified by progesterone levels as designated in Fig. 1.

**Figure 1.**
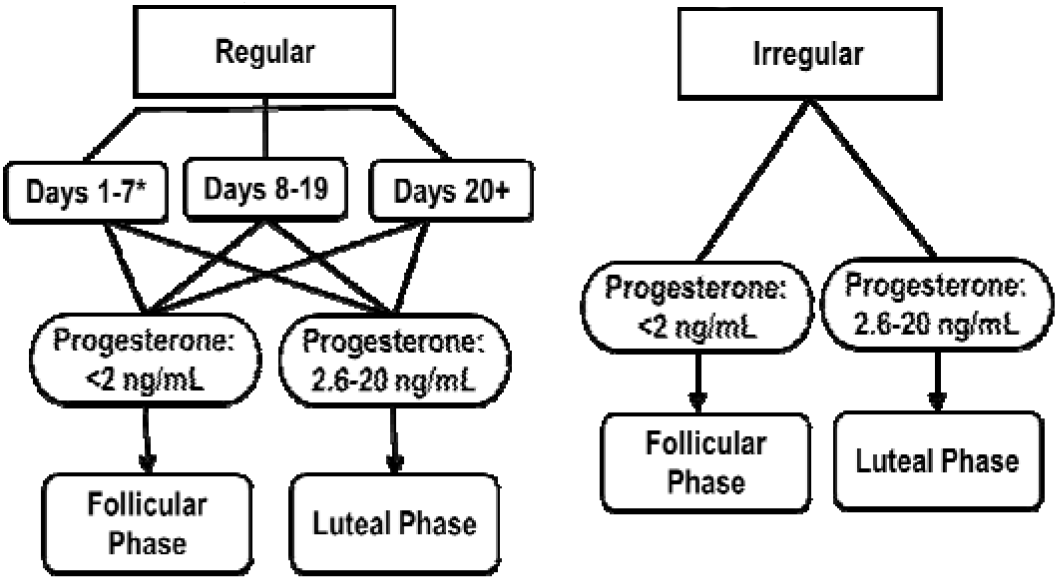
Determining menstrual cycle phase in non-hormonal contraceptive users. *Day 1 is first day of bleeding; Day of cycle is based on retrospective and prospective, when available, self-reported tracking of menstrual cycles.

### Hormonal contraceptive user classification

By definition, regular hormonal contraceptive users do not have typical MCs, defined by the progesterone surge that occurs after ovulation^17^. Although some hormonal contraceptive users bleed, this is not synonymous with menstruation^18^. Studies demonstrate that females using various types of oral contraceptive pills typically have suppressed progesterone levels across the pill pack, ranging from .30-.76 ng/mL (SD ~ .5 ng/mL)^19,20^. In general, hormones, including serum progesterone and estrogen, remain consistent and suppressed, relative to non-hormonal contraceptive users, across the pill pack^21^. Therefore, hormonal contraceptive users using an oral contraceptive pill, patch or NuVA Ring were classified into “active-HC” (e.g., when taking non-placebo, active hormone pills or when NuvaRing was inserted) or “inactive-HC” (e.g., when taking placebo, inactive hormone pills or when NuvaRing was not inserted) (Fig. 2).

**Figure 2.**
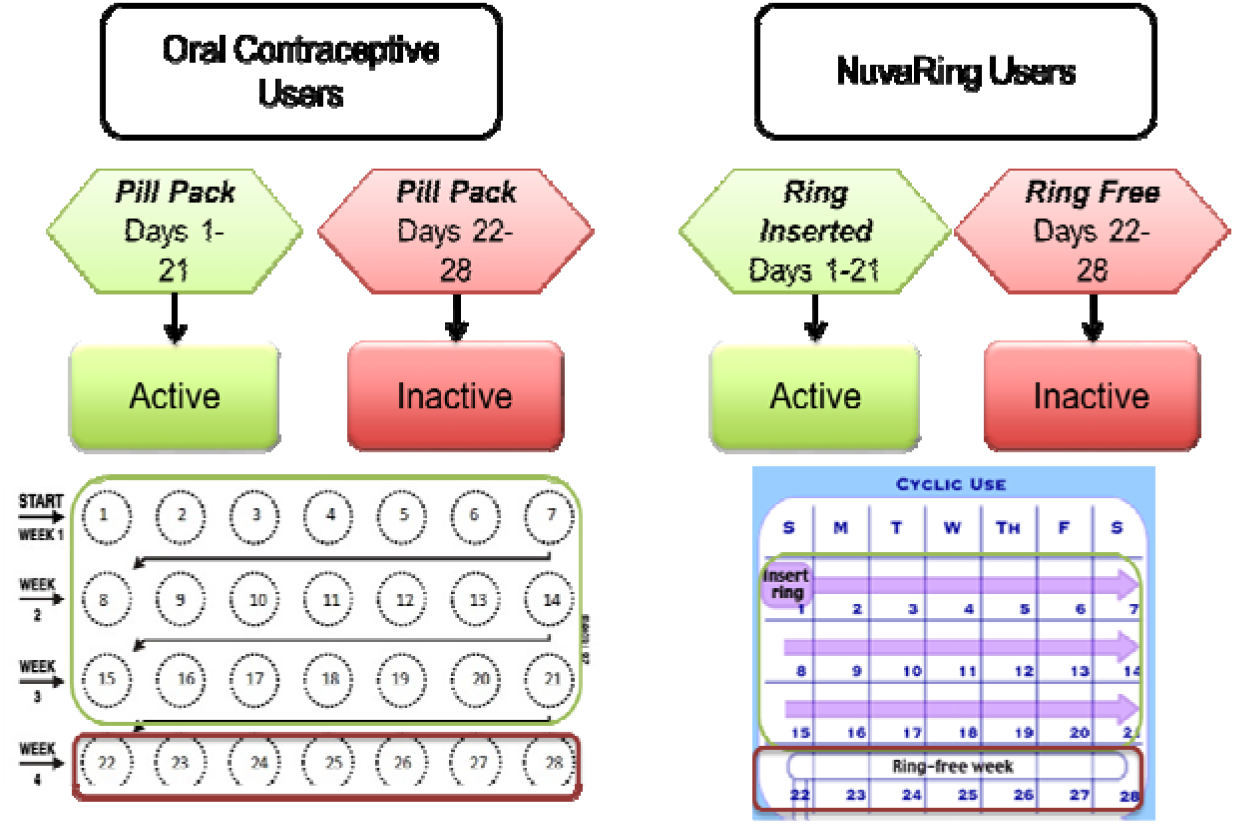
Determining phase in hormonal contraceptive users. Active phases are illustrated in **green** and inactive phases are illustrated in **red**.

### Matching control participants based on MC phase among non-hormonal contraceptive users and active vs. inactive phase among hormonal contraceptive users

Among non-hormonal contraceptive users, we selected demographically matched controls who had three cycles of self-reported menstrual cycling tracking data (electronically reported first day of period). If the mTBI athlete was determined to be in the follicular phase, we invited the matched control to attend the study visit in days 1-7 of MC. Follicular phase was confirmed with progesterone testing on the day of the study visit. Among hormonal contraceptive users, we selected matched controls who are on the same form on hormonal contraception (e.g., monophasic hormonal contraceptive pill, triphasic hormonal contraceptive pill, NuvaRing, or implant). Control participants were required to be on their form of contraception for at least 6 months. Control participants were matched by active vs. inactive phase and then invited for their study visit within 2 days of the matched mTBI participants’ pill pack day or NuvaRing day.

### Data Analysis

Descriptive analyses of demographics and sample characteristics between groups (mTBI and CON) were conducted using Student’s *t*-tests for continuous variables and Chi-Squared tests for categorical variables (α = 0.05). Descriptive analyses of frequencies and percentages of number of athletes in using and not using hormonal contraceptives and those in the follicular and luteal MC phase were conducted. For the non-hormonal contraceptive users, we calculated both percent agreement and a kappa statistic for identification of MC phase in order to inform accuracy. We used the following equation to calculate κ = (p_o_ − p_e_) / (1 − p_e_), where p_o_ is the relative observed agreement among self-report and progesterone methods of identifying MC phase, and p_e_ is the hypothetical probability of chance agreement, using the observed data to calculate the probabilities of each method randomly seeing each category (i.e., follicular or luteal).

## Results

### Sample Characteristics

Table 1 summarizes the sample demographics and sample characteristics (N=30). The mTBI and CON groups (n=15 each) were similar in age, race, ethnicity, socioeconomic status, pre-morbid IQ, and history of depression and anxiety. The only factor that differed between the two groups was self-reported history of previous mTBIs (p=.003). The number of self-reported previous concussions was greater for the mTBI (mean .53, stdev .64) than for the CON group (mean 0, stdev 0).

**Table 1.** Sample Demographic Information

| Table 1. |  |  |  |
| --- | --- | --- | --- |
| *Sample Demographic Information* |  |  |  |
|  | Group |  | p |
|  | mTBI (n=15) | CON (n=15) |  |
| Age [M(SD)] | 20.19 (1.13) | 20.20 (1.53) | ns |
| Race (% Caucasian) | 87% (13/15) | 93% (14/15) | ns |
| Ethnicity (% Hispanic or Latino) | 13% (2/15) | 13% (2/15) | ns |
| Years of Education [M(SD)] | 13.07 (1.22) | 13.07 (1.01) | ns |
| WTAR Standard Score [M(SD)] | 113.53 (11.04) | 118.00 (6.47) | ns |
| Hx of Anxiety | 0% (0/15) | 0% (0/15) | ns |
| Hx of Depression | 7% (1/15) | 0% (0/15) | ns |
| Number of previous concussions | .53 (.64) | 0 (0) | .003 |
| Regular MC pattern during ages 18-22 | 87% (13/15) | 67% (10/15) |  |
| <u>Average MC length over past year:</u> |  |  |  |
| • 21-25 days | 33.3% (5/15) | 6.7% (1/15) |  |
| • 26-32 days | 53.3% (8/15) | 53.3% (8/15) |  |
| • 36-90 days | 6.7% (1/15) | 6.7% (1/15) |  |
| • More than 90 days | 6.7% (1/15) | 0% (0/15) |  |
| • Too variable to say | 0% (0/15) | 20% (3/15) |  |
| • Don't know | 0% (0/15) | 6.7% (1/15) |  |

The Phenx Menstrual Cycle history questionnaire provided general self-report information about the participants MC (Table 1). Most females in both the mTBI (87%) and CON (67%) groups self-reported regular menstrual cycles occurring regularly (approximately once per month). As such, only 1 participant, in each group self-reported MC between 26-90 days in length. There was only 1 participant, who was in the mTBI group, who self-reported a MC longer than 90 days. Three CON participants reported MCs that were too variable to say, and 1 CON participant reported that they did not know about their MC length. Notably, no participants identified not having a period or that they were anovulatory.

The females on HC all self-reported being on these medications at least 6 months prior to completing assessments.

### Menstrual Cycle (MC) Phase Matching and Characteristics

Table 2 summarizes matching of female athletes with mTBI (n=15) to healthy control athletes without a recent history of mTBI (n=15) based on MC phase and hormonal contraceptive use at time of assessment (3-10 days post-mTBI for the mTBI group).

**Table 2.** Matching mTBI to Control (CON) Female Athletes on Menstrual Cycle Phase and Hormonal Contraceptive Use

|  | Group | Progesterone (ng/mL) | Hormonal Contraceptive Type | Menstrual Cycle Phase | Hormonal Contraceptive Phase |
| --- | --- | --- | --- | --- | --- |
| **Non-Hormonal Contraceptive Users (with Natural Menstrual Cycles)** |  |  |  |  |  |
| Pair 1 | mTBI | 0.26 |  | Follicular |  |
|  | CON | 0.30 |  | Follicular |  |
| Pair 2 | mTBI | 0.27 |  | Follicular |  |
|  | CON | 0.08 |  | Follicular |  |
| Pair 3 | mTBI | 0.68 |  | Follicular |  |
|  | CON | 0.24 |  | Follicular |  |
| Pair 4 | mTBI | 0.31 |  | Follicular |  |
|  | CON | 0.72 |  | Follicular |  |
| Pair 5 | mTBI | 0.44 |  | Follicular |  |
|  | CON | 0.40 |  | Follicular |  |
| Pair 6 | mTBI | 0.31 |  | Follicular |  |
|  | CON | 0.40 |  | Follicular |  |
| Pair 7 | mTBI | 12.23 |  | Luteal |  |
|  | CON | 18.06 |  | Luteal |  |
| Pair 8 | mTBI | 0.35 |  | Follicular |  |
|  | CON | 0.25 |  | Follicular |  |
| **Hormonal Contraceptive Users** |  |  |  |  |  |
| Pair 9 | mTBI | 0.49 | Implant |  | Active |
|  | CON | 0.28 | OCP |  | Active |
| Pair 10 | mTBI | 0.53 | OCP |  | Active |
|  | CON | 0.39 | OCP |  | Active |
| Pair 11 | mTBI | 0.80 | OCP |  | Inactive |
|  | CON | 0.32 | OCP |  | Inactive |
| Pair 12 | mTBI | 0.38 | OCP |  | Inactive |
|  | CON | 0.41 | OCP |  | Inactive |
| Pair 13 | mTBI | 1.21 | Nuva Ring |  | Active |
|  | CON | 1.39 | OCP |  | Active |
| Pair 14 | mTBI | 0.2 | OCP |  | Inactive |
|  | CON | 1.44 | OCP |  | Inactive |
| Pair 15 | mTBI | 0.32 | OCP |  | Inactive |
|  | CON | 0.47 | OCP |  | Inactive |

CON, control; OCP, Oral Contraceptive; NA, not applicable

Table 3 summarizes cross-classification accuracy between self-report and progesterone methods for identification of MC phase among the 16 non-hormonal contraceptive users. Six females self-reported the day of their MC between days 8-19 (in-between or possible ovulation) where it is difficult to reliably predict MC phase based on self-report alone. Thus, we did not calculate percent agreement or κ utilizing these athletes. We calculated 50% (5/10) agreement between self-report and progesterone level and κ = 0.138, which is considered poor^22^.

**Table 3.**
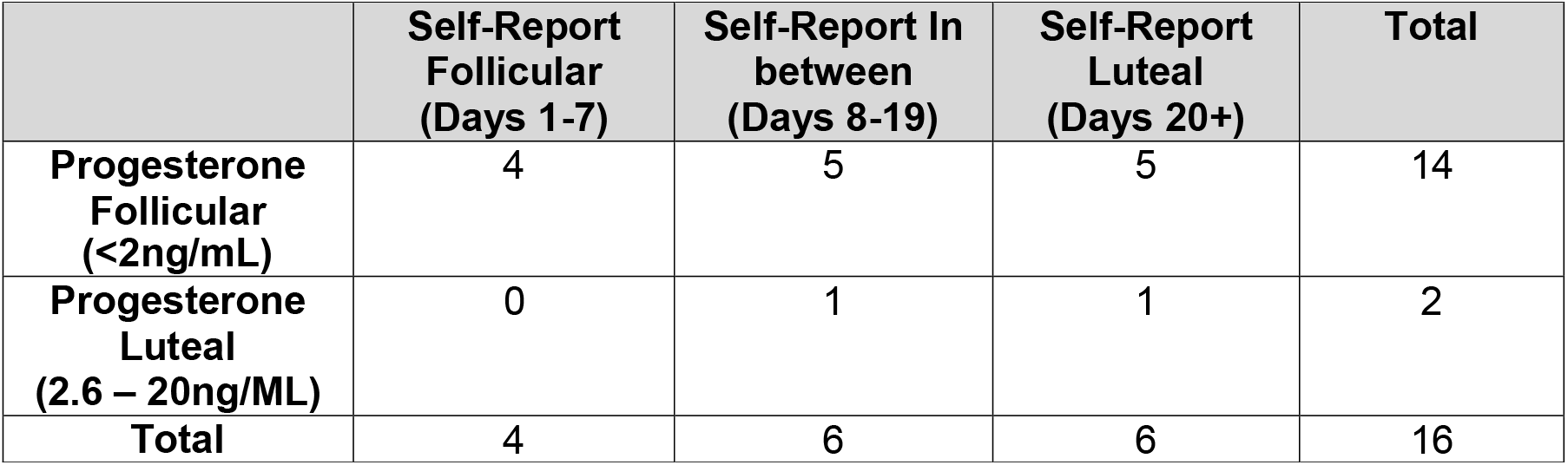
Cross-classification table for non-hormonal contraceptive users (n=16)

For the 14 hormonal contraceptive users (Table 4), obtaining progesterone level data allowed us to confirm compliance with medication, as progesterone levels were less than 0.9ng/mL^20^ (average = .58ng/mL).

**Table 4.**
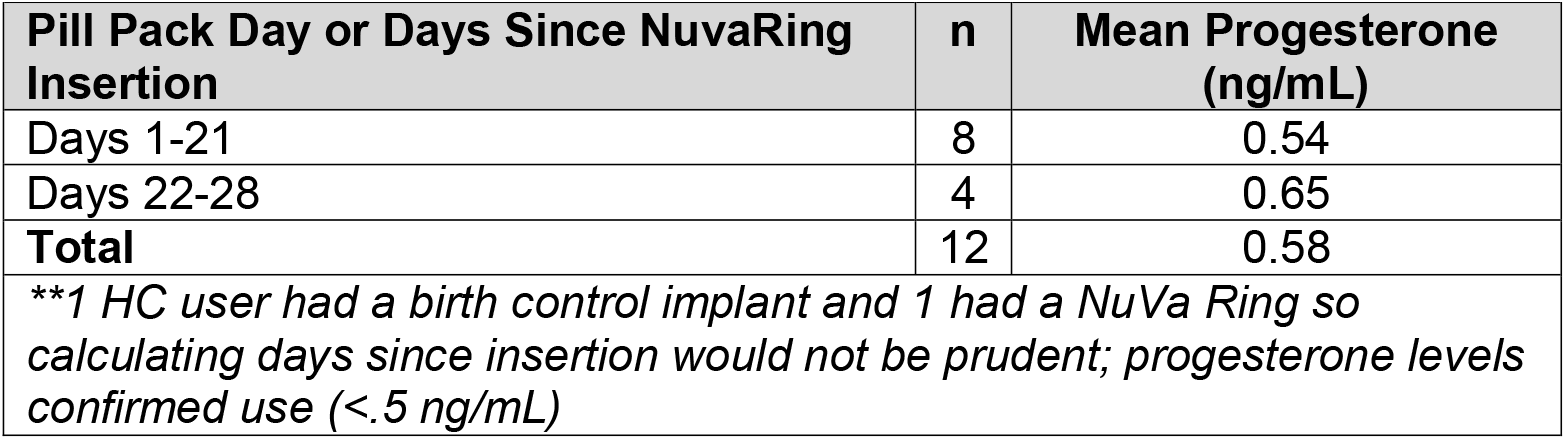
Hormonal classification of hormonal contraceptive users (n=14).**

## Discussion

This novel study addresses the unmet need of testing methods of controlling for the female-specific factor of menstrual cycle (MC) phase in prospective translational research. This study was conducted in a sample of female collegiate athletes who recently experienced a mTBI. This is of great importance given that females are often understudied, grouped together with males, and female-specific factors are often not accounted for when interpreting study outcomes. In our study, 50% of the sample were hormonal contraceptive (HC) users and 50% were not. Among non-HC users, self-report of MC phase was 50% accurate for predicting and matching controls when compared to progesterone levels with κ = 0.138, which is considered poor, indicating that self-report measures can be markedly improved by the addition of objective progesterone level data. Among HC users, progesterone levels confirmed HC medication compliance.

Among the non-HC users, fourteen female pairs were in the follicular and two were in the luteal phase. Notably, 6 females self-reported the day of their MC between days 8-19 where it is difficult to reliably predict MC phase based on self-report alone. During this time 48-hour period of ovulation could be occurring and this cannot be determined without objective, hormonal confirmation. Thus, during this phase, we suggest relying upon objective hormone levels. We also noted that self-report was less reliable among the mTBI than the CON athletes. This is not surprising given that symptoms of mTBI involve cognitive and memory impairments. In addition, a methodological feature of our study was such that for the CON group we were not only able to gather retrospective self-report MC phase data but also prospective self-report MC phase data, which is less prone to report bias. Thus, we suggest that among clinical samples with conditions that involve cognitive impairment, utilizing objective hormone measures to confirm MC phase would be extremely valuable.

Among HC users, all users had progesterone levels that confirmed medication compliance. However, there was a large amount of variability and overlap in progesterone levels between the active and in-active HC phase. This could be due to the fact that while we did match on the type of HC, we did not match on the exact medication and synthetic progesterone levels in each medication can be different. In addition, there can be variability in the time of day the medications are taken relative to when the blood draw for progesterone was taken, as well as individual metabolic differences that could account for this variability.

This study must also be described in terms of its limitations. We had a limited, pilot sample size of 30 participants including a single site of highly functional and intelligent collegiate students, who were also predominantly Caucasian. This indicates that future studies including multiple sites with increased racial and ethnic diversity are warranted. In addition, while progesterone is the best objective, hormonal indicator of MC phase. There are circumstances in which accuracy of determining MC phase could be strengthened by also including luteinizing hormone (LH) surge or estradiol measures. The addition of LH surge measures would assist in identification of the 48-hour ovulation period in which females are not clearly in the follicular or luteal phase. Estradiol measures, may also help provide another reference point, which would be helpful when the progesterone levels are between 2-2.6ng/mL Finally, serial hormonal measurements collected longitudinally over time may also provide further accuracy in determining MC phase for future work that may not have as many time or resource constraints.

In conclusion, we have provided algorithms that can be utilized to determine MC phase among female adolescent athletes. These algorithms were developed and tested in a sample of HC users and non-users; thus, increasing generalizability. Conducting this study among female athletes 3-10 days post-mTBI demonstrates the feasibility of utilizing this method in time-sensitive prospective translational research studies in symptomatic, clinical samples. This is an important contribution, given the National Institute of Health mandate that not only females and males be studied but sex as a biological variable must be addressed in all funded studies.

